# Predictive model identifies strategies to enhance TSP1-mediated apoptosis signaling

**DOI:** 10.1101/188003

**Authors:** Qianhui Wu, Stacey D. Finley

## Abstract

This study explores strategies to enhance thrombospondin-1 (TSP1) induced apoptosis in endothelial cells, an important function that contributes to TSP1’s anti-angiogenic effect. We established a mathematical model of TSP1-mediated intracellular signaling via the CD36 receptor. This model was used to investigate the effects of several approaches to perturb the TSP1-CD36 signaling network. Model simulations predict the population-based response to strategies to enhance TSP1-mediated apoptosis, such as downregulating the apoptosis inhibitor XIAP and inhibiting phosphatase activity. The model also postulates a new mechanism of low dosage doxorubicin treatment in combination with TSP1 stimulation. Using computational analysis, we predict which cells will undergo apoptosis, based on the initial intracellular concentrations of particular signaling species. Overall, the modeling framework predicts molecular strategies that increase TSP1-mediated apoptosis, which is useful in many disease settings.

## Introduction

Angiogenesis, the formation of new capillaries from pre-existing blood vessels, plays a critical role in tumor progression. Angiogenesis enables the tumor to generate its own blood supply and obtain oxygen and nutrition from the microenvironment. This process is regulated by a dynamic interplay between the angiogenic promoters, such as vascular endothelial growth factor (VEGF) and fibroblast growth factor (FGF), as well as angiogenic inhibitors, such as thrombospondin-1 (TSP1) [1–5].

Due to its importance in tumor development, invasion, and metastasis, angiogenesis has become a prominent target for cancer therapies. In addition to strategies targeting pro-angiogenic species, such as inhibiting VEGF signaling using antibodies and tyrosine kinase inhibitors, anti-angiogenic species hold promise in reducing tumor angiogenesis. TSP1 is a well-known, potent endogenous angiogenesis inhibitor. TSP1 expression is lost in multiple cancer types; however, its re-expression can delay cancer progression, promote tumor cell apoptosis, and decrease microvascular density. For these reasons, it has been of interest to mimic TSP1’s functions in regulating angiogenesis [3,7–10].

TSP1 is a multifunctional matricellular protein that acts to inhibit angiogenesis in multiple ways [2,11,12], which include altering the availability of pro-angiogenic factors and promoting anti-angiogenic signaling through its receptors CD36 and CD47. Several studies have shown that TSP1 mediates its anti-proliferative and pro-apoptotic effects in a highly specific manner on endothelial cells. TSP1 primarily promotes these effects by binding to the CD36 receptor [3,13,14], which is associated with capillary blood vessel regression [11,13,15,16]. TSP1 interaction with CD36 leads to recruitment of the Src-related kinase Fyn, activation of p38MAPK, and processing of caspase-3, a vital protease that mediates apoptosis [13,16,17]. TSP1-CD36 signaling also causes transcriptional activation of Fas ligand (FasL), a death ligand that also promotes pro-apoptotic signaling, ultimately inhibiting angiogenesis. This apoptosis pathway is further enhanced as pro-angiogenic factors induce increased levels of Fas receptors, sensitizing the cells to FasL [17].

Unfortunately, therapies that mimic TSP1 activity have not demonstrated definitive clinical efficacy. For example, ABT-510, a TSP1 peptide mimetic that binds to CD36, was previously tested in a Phase II study in 2007 for treatment of metastatic melanoma. However, the drug failed to reach its primary endpoint (18-week treatment failure rate), resulting in termination of the study [18]. ABT-510 also showed little clinical effect in a Phase II trial for advanced renal cell carcinoma [19]. These disappointing results indicate that there is a need to better understand the effects of anti-angiogenic agents and develop effective treatment strategies. This requires a detailed and quantitative understanding of the dynamic concentrations of the factors involved in angiogenesis signaling.

Computational systems biology offers powerful tools for studying complex biological processes that involve a large number of molecular species and signaling reactions that occur on multiple time‐ and spatial-scales. Systems biology aims to study how individual components of biological systems give rise to the function and behavior of the system [20]. Additionally, computational modeling aids in the development of therapeutic strategies that specifically target tumor angiogenesis to optimally inhibit tumor progression, complementing pre-clinical and clinical angiogenesis research [21].

Substantial research has focused on the pro-angiogenic factors and their extracellular interactions [22–24]. However, a consideration of the intracellular mechanisms of anti-angiogenic factors is also needed in order to fully understand the dynamics of the signaling networks regulated by angiogenesis promoters and inhibitors. In this study, we focus on TSP1-mediated apoptosis signaling through the CD36 receptor. Although some aspects of the TSP1-CD36 pathway have been studied experimentally, the signaling network has not been quantitatively and systematically analyzed. We constructed the first computational model that describes the intracellular signaling network induced by TSP1-CD36 binding in endothelial cells, a complex network comprised of biochemical reactions that lead to cell apoptosis. We apply the model to predict the effects of modulating protein expression and enzyme activity on apoptosis signaling. The model quantifies the effects of these perturbations and predicts promising targets, both in terms of the averaged response of a population of endothelial cells and individual cells within the population. Thus, the model is a quantitative framework to predict strategies to enhance TSP1-mediated apoptosis. Ultimately, the model can be used to identify novel pharmacologic targets and optimize therapeutic strategies that promote apoptosis and, subsequently, inhibit angiogenesis.

## Methods

### Mathematical model

We constructed a computational model of TSP1-mediated apoptosis signaling via the CD36 receptor in endothelial cells. The molecular interactions depicted in Figure 1 were translated into biochemical reaction equations, with the assumption that the reactions follow well-established kinetic laws, including mass-action or Michaelis-Menten kinetics (Table S1). A system of nonlinear ordinary differential equations (ODEs) was formulated to describe the rate of change of the species’ concentration. The model is comprised of 53 ODEs to predict the concentrations of the 53 species in the signaling network over time. The Simbiology toolbox (MATLAB) was used to implement the biochemical reaction equations, and the MATLAB stiff solver ODE15s was used the numerically solve the system of ODEs. The model file is provided in the Supplementary Materials.

**Figure 1.**
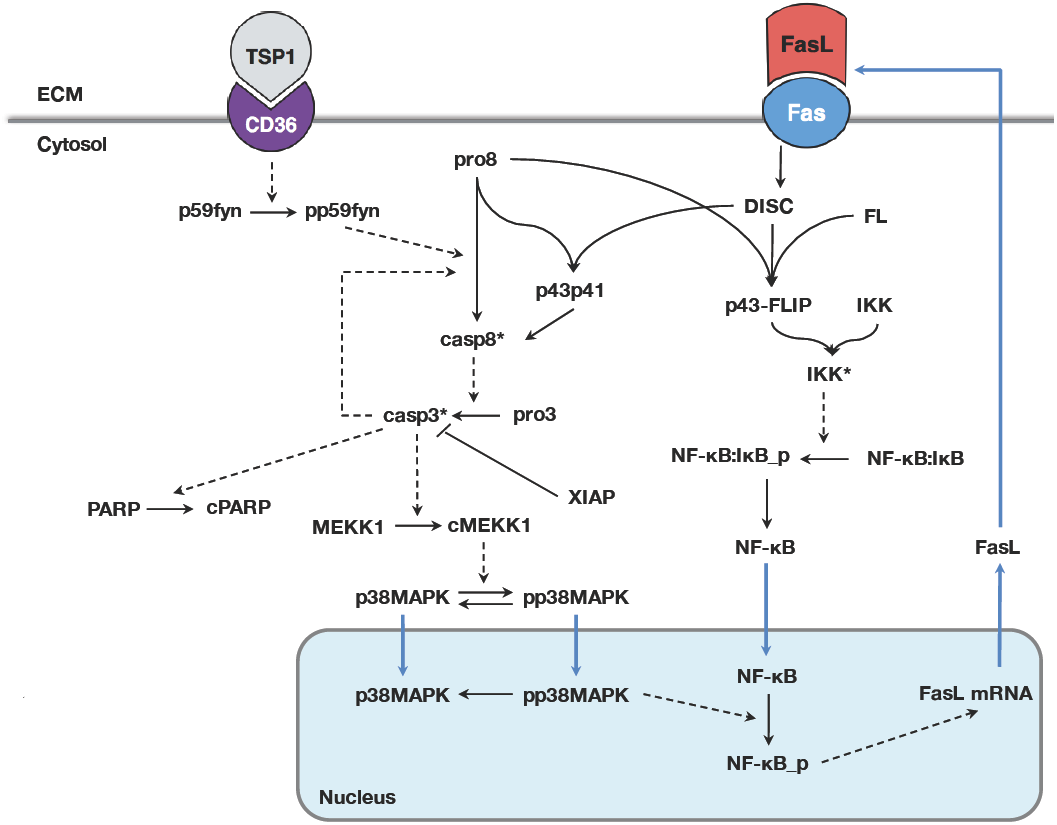
Model schematic of TSP1-mediated apoptosis signaling. TSP1 binding to the CD36 receptor recruits p59fyn, which induces activation of the caspase-3 cascade. The kinase p38MAPK is subsequently phosphorylated and translocated to the nucleus. NF-kB translocates into the nucleus and is activated in presence of phosphorylated p38MAPK. This leads to transcriptional activation of FasL. FasL protein binds to its receptor Fas, forming the DISC complex, which binds to c-FLIP (FL) and procaspase-8 (pro8) to form the p43-FLIP complex. This complex activates IKK, which releases NF-kB from its inhibitor IkB. Blue arrows indicate transport reactions.

Solving the set of ODEs with the baseline initial conditions provides the averaged response of a population of cells. Additionally, we account for heterogeneity in a population of cells by solving the ODE model 2,000 times, each with a different set of initial conditions. We refer to this as the “population-based model”.

### Cytosolic and nuclear compartments

The model is comprised of two compartments, cytosolic and nuclear, both assumed to be well mixed. Specific molecules, such as NF-kB, may move from one compartment to another at a defined translocation rate. The volume of the nuclear compartment is estimated to be 14.32% of the cytosolic compartment [24], and the concentrations of species transported between the two compartments is converted using this ratio.

### Initial protein concentrations

The initial conditions used in the model are given in Table S2. Very few references for initial concentrations of proteins are available. Therefore, we adapted values from a previously published model [25] and adjusted the initial concentrations of several species in order for the model to match experimental measurements. For the CD36 and Fas receptors, we used flow cytometry to quantify the average numbers of receptors on cultured human microvascular endothelial cells (data not shown; similar to previous work [26], and converted the receptor numbers to concentrations using the total cell volume of 1 picoliter [27].

### Rate constants

All baseline model parameter values are listed in Table S3. *Production of soluble species.* The basal rate at which each species is synthesized (*K*_*syn_all*_) is set to be 10”^4^ min”^1^, with the exception of FasL, procaspase-8, and procaspase-3, whose production rates are described below.

The model accounts for FasL production mediated by TSP1, and we described the production of FasL mRNA production (DNA transcription) using Michaelis-Menten kinetics:

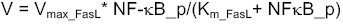

where *V*_*maxFasL*_ and *K*_*mFasL*_ and the Michaelis-Menten kinetic rate constants for FasL mRNA production, and *NF-kB_p* is the activated transcriptional factor that catalyzes this process. The molecular details involved in FasL protein production encompass the mRNA translocation and translation, and protein secretion. The rates involved in these reactions are not readily available in published literature. Therefore, we estimated the values in model fitting in order to match experimental data.

The synthesis rate of procaspase-8 and procaspase-3 were assumed to be dependent on the concentration of DISC present in the system, as a partial effect of Fas ligation. The synthesis rate is described as:

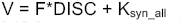

where *F* is a hand-tuned coefficient, *DISC* is the complex formed by FasL binding to Fas, and *K_sy_n_aii* is the basal level synthesis rate assigned to all the other species except for FasL.

*Protein degradation.* Protein species are assumed to be degraded at the same rate, 10∼^3^ μM/min, unless there was a degradation rate available in the literature or from a previous model. This allows the system to balance and reach steady-state in the absence of TSP1 stimulation. The degradation rates of caspase-8, caspase-3, the p43:FLIP:IKK_a complex, and cytosolic NF-kB have unique values adapted from previous modeling work by Neumann *et al.* [25].

*Receptor-ligand interactions.* The affinity of TSP1 and its receptor CD36 has been measured experimentally: the *K_d_* value is 230 nM [28]. We assume that FasL binds to Fas with an affinity of 0.4 nM. In all cases, the dissociation rate for the receptors is 1.2x10”^2^ min”^1^. Receptors are internalized and inserted at the cell membrane such that the total number of receptors (ligated plus unbound) is constant.

*FasL cascade.* The model includes DISC formation upon FasL binding with Fas, and the downstream caspase-8 and NF-kB activation reactions. The molecular details were adapted from the model established by Neumann *et al.* [25]. We altered this portion of their model by adding reversible binding reactions to ensure the reaction network is consistent with the other parts of our model. We tuned the universal dissociation rate *K_off_* to be 1.2x10”^2^ min”^1^ to match the data presented in their paper. The simulations of the implemented minimal model are shown in Supplemental Figure S1.

### Sensitivity analysis

There is limited quantitative experimental data available to specify the values of the kinetic parameters. However, the parameters must be set to appropriate values in order for the model to generate reliable predictions. We first used sensitivity analysis to reduce the number of parameters to be estimated. Specifically, to identify the influential kinetic parameters before each step of model fitting, we conducted global sensitivity analysis using the extended Fourier Amplitude Sensitivity Test (eFAST) method [29], as we have done in previous work [23,24]. All inputs were allowed to vary simultaneously one order of magnitude above and below the baseline value, and the effects of multiple inputs on the model outputs of individual inputs were quantified. This sensitivity analysis was also used to determine the effects of initial protein concentrations to inform perturbation simulations (Figure S2).

### Quantification of experimental data

The Western blots were analyzed using ImageJ (https://imagej.nih.gov). The local background from bands was subtracted and their intensity was quantified. The intensities of subject species were normalized to the corresponding control band intensities.

### Parameter estimation

The estimation of the kinetic parameters was achieved using the “lsqnonlin” function in MATLAB, as done in our previous work [24,42]. This algorithm solves the non-linear least squares problem using the trust-region-reflective optimization algorithm, minimizing the weighted sum of the squared residuals (WSSR). The minimization is subject to the upper and lower bounds of the free parameters. One hundred runs were performed in each fitting step, and a global sensitivity analysis was performed with the best fit parameter values (the parameters that produce the lowest WSSR). The step-wise iteration was repeated four times to ensure fine-tuning of the parameter values. Parameter values used in the implemented model are from the best fit (lowest WSSR) from the last step. We also report the mean and standard deviation of the estimated parameter values in Table S3 and Supplemental Figure S3.

### Definition of apoptotic cells

Cleaved poly(ADP-ribose) polymerase (cPARP) is the output of the model used as an indicator of apoptosis, since loss of intact PARP results in failure to repair DNA damage. Our model simulations show that the dynamics of cPARP follows a switch-like action; however, the range of cPARP varies widely depending on the initial concentration of PARP. Previous study [32] has shown that low DXR dosage with 10 nM TSP1 stimulation resulted in approximately 50% of the cells becoming apoptotic in 24 hours. Therefore, we simulated this treatment condition using the population-based model, and determined the cPARP concentration that results in 50% cell apoptosis. We then use this concentration, 1.05 nM, as the defined threshold that needs to be reached for cell apoptosis to occur. Thus, the definition of which cells are apoptotic is based on literature evidence.

### Simulated perturbations to TSP1-mediated apoptosis

We apply the model to simulate seven specific perturbations to the intracellular signaling network, to find strategies that enhance apoptosis signaling. The abbreviations listed in parentheses are also used in the results figures. Generally, perturbations are simulated in the ODE model by adjusting the baseline initial conditions (Table S2) and parameter values (Table S3).

1. XIAP downregulation (“XIAP”): XIAP downregulation can promote the apoptotic signaling [33]. We simulated this effect by reducing XIAP concentration to 0.5-fold of the baseline value.
2. Low dosage doxorubicin treatment (“DXR”): Experimental studies [27,28] have shown that a low dose of doxorubicin upregulates the expression of Fas receptor and other protein species. We simulated this effect by increasing the initial Fas receptor level by 3‐ fold and *K*_*syn_all*_ by 10-fold.
3. Phophatase inhibition (“Ptase”): Studies have shown [31,32] that inhibiting MAPK phosphatase (MKP) activity can promote apoptosis signaling. We simulated this effect by decreasing the association rate *(K_on___dephos_)* of the phosphatase with phosphorylated p38MAPK (pp38) and the dephosphorylation rate *(K_dephos_)* by 10-fold.
4. Kinase promotor (“Kp”): Literature suggest that the tumor microenvironment likely upregulates many kinase’s activity in the tumor-related endothelial cells [37]-[39]. We simulated the kinase promoter by increasing the phosphorylation rates of p59fyn, P38MAPK, and IkB by 10-fold.
5. Procaspase-3 upregulation (“pro3”): Global sensitivity analysis (Figure S2) and baseline model simulations (Figure 3) indicates that upregulation of procaspase-3 increases the cPARP leveluponTSP1stimulation. We simulated the effect of procaspase-3 upregulation by increasing procaspase-3 concentration by 3-fold.
6. Fas upregulation(“Fas”):ExperimentalstudybyQuesada*et al.*suggest that upregulation of the receptor Fas promotes TSP1-induced apoptosis [32]. We simulated the effect of Fas upregulation by increasing Fas concentration by 3-fold.
7. Translocation rate increase (“Ktrsp”): Based on the structure of the signaling network, we hypothesizedthatincreasingthecytoplasm-to-nucleustransport would enhance apoptosis. We simulated the effect of faster cytosol-to-nucleus translocationby increasing the translocation rate (K,_rsp_) by 10-fold.

### Analysis of sensitivity and specificity of a cPARP-based classifier

Here, we consider binary classification of the cells’ response to TSP1 stimulation based on cPARP levels at 24 hours: apoptotic or non-apoptotic. For a binary classification system such as this, there are four possible predicted outcomes for a given cPARP cutoff value: *true positive,* a cell predicted to be apoptotic is actually apoptotic; *false positive,* a cell predicted to be apoptotic is actually non-apoptotic; *true negative,* the predicted and actual response are both non-apoptotic; *false negative,* a cell predicted to be non-apoptotic is actually apoptotic. This analysis determines the fraction of positives predicted correctly (sensitivity or the true positive rate) and the fraction of true negatives predicted (specificity or the true negative rate) for different cutoff values of cPARP.

To evaluate tradeoffs between sensitivity and specificity, we constructed a receiver operator characteristic (ROC) curve for cPARP. The ROC curve plots the true positive rate versus the false positive rate (1-specificity). An ideal input maximizes true positives, with minimal false positives (i.e., the (0,1) point on the ROC graph). An ROC curve that lies on the 45-degree angle line indicates that the input does not classify the output any better than a random guess, where the area under the ROC curve (AUC) is 0.5. Thus, having an AUC value significantly greater than 0.5 indicates that the input can be used to classify the data. We performed the ROC analysis using the custom *“roc”* function in MATLAB.

## Results

### Model training and validation

We constructed a model of the signaling network of TSP1-mediated apoptosis in endothelial cells based on literature evidence. TSP1 binds to CD36, activating caspase-3, the core executioner protease. Caspase-3 promotes apoptosis by cleaving PARP in endothelial cells. Activation of caspase-3 also mediates intracellular signaling leading to the production of FasL, a death ligand that binds to its receptor Fas on endothelial cells and further promotes apoptosis through activation of caspase-3 [13,16,18]. The signaling network, illustrated in Figure 1, includes several important feedback loops involved in TSP1-mediated apoptosis, including the caspase cascade (caspase-3 activates its activator, caspase-8) and Fas signaling (TSP-1 promotes the production of Fas, which also activates caspase-3). We implemented the signaling network mathematically to generate an ODE model, assuming that the reactions follow mass-action or Michaelis-Menten kinetics rate laws.

The model was trained using quantitative experimental data and validated with an independent set of measurements. We extracted experimental data from the literature in order to calibrate the model and estimate the kinetic parameters. Specifically, the fold-changes in the caspase-3 activity and the levels of two intracellular species (p59fyn and p38MAPK) upon TSP1 stimulation were quantified from Western blot data and used to train the ODE model.

We used a step-wise strategy comprised of global sensitivity analysis and parameter estimation to ensure that the model could match the training data (see Methods). As a result of this approach, we obtained 12 sets of parameters that enable the model to closely reproduce the training data (Figure 2A-D).

**Figure 2.**
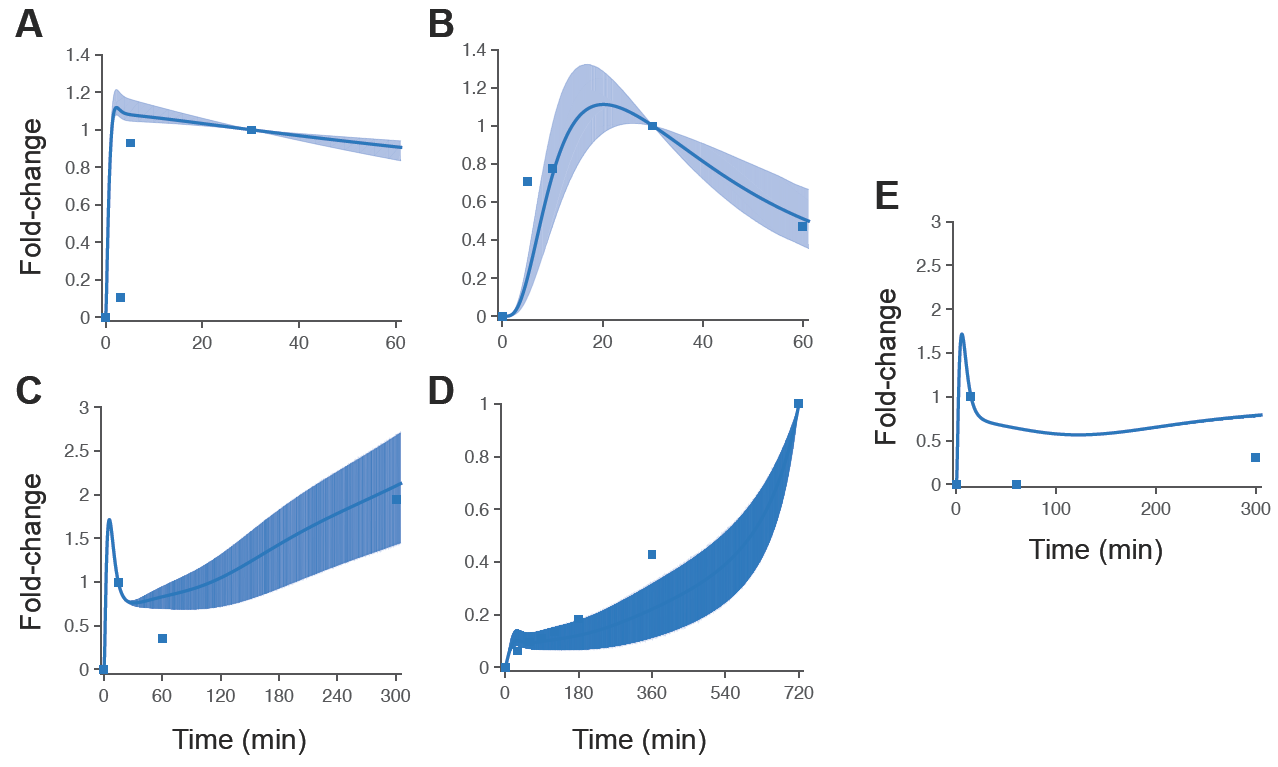
Model training and validation. The ODE model was trained to match experimental measurements of activated species in the TSP1-mediated apoptosis signaling pathway. A) pp59fyn [12]; B) pp38MAPK [12]; C) caspase-3 activity [12]; and D) caspase-3 activity [15]. E) An independent set of data for caspase-3 activity under the condition of p38MAPK inhibition [12] was used to validate the model prediction. Solid line: mean of 12 best fits. Shaded area: 95% confidence interval. Squares: experimental data.

After fitting the model to the experimental data, we used a separate set of measurements to validate the model predictions. Here, we applied the trained model to predict the dynamics of caspase-3 activity when p38MAPK is inhibited, mimicking an experimental study from Jimenez *et al.* [12]. This inhibitory effect on p38MAPK is simulated by setting the phosphorylation rate of NF-kB by active p38MAPK (pp38MAPK) to be zero. The model qualitatively matches this independent set of data (Figure 2E), where caspase-3 activity is reduced at 300 minutes, compared to the case without p38 MAPK inhibition (Figure 2C). Overall, the model fitting and validation produces a trained model that generates reliable predictions related to the dynamics of TSP1 simulation.

### Altering the concentrations of intracellular signaling species influences the apoptotic response

We first applied the trained and validated model to investigate the effects of varying the concentrations of cell surface receptors and intracellular signaling species, in combination with different TSP1 stimulation levels. In this study, we specifically focus on predicting the concentration of cleaved PARP (cPARP) as an indicator of cell apoptosis. Caspase-3 promotes apoptosis by cleaving PARP, and cleavage of PARP by caspases is considered a hallmark of apoptosis [40]. Sensitivity analysis revealed that the concentrations of procaspase-3, XIAP, and PARP most significantly influence the cPARP level throughout the simulated time course (Figure S2). This analysis suggests that varying the concentrations of those intracellular species can impact on TSP1-mediated apoptosis signaling. We also hypothesized that increasing the receptor’s availability (i.e., increasing the receptor: ligand ratio) can amplify the signaling induced by ligand-receptor binding. Therefore, we ran the model and individually altered the expression level (initial conditions) of the CD36 or Fas receptors, or intracellular species procaspase-3, XIAP, and PARP, by 10-fold above and below the baseline values. We applied this relatively large alteration in the protein expression levels to explore the extent of changes in the model output. The initial conditions were varied for each of the 12 fitted parameter sets, and we compared the cPARP level at various time points for each case.

Across the simulated time points, there is a dose-dependent response to TSP1, where increasing the concentration of TSP1 increases the predicted cPARP concentration. Interestingly, altering the expression levels of the CD36 or Fas receptors does not affect the cPARP level, compared to the baseline model (Figure 3A-B). This result holds true for all TSP1 concentrations investigated. In contrast, increasing the amount of procaspase-3, the unprocessed form of caspase-3, by 10-fold leads to increased cPARP level at every simulated time point, as compared to the baseline model (Figure 3C, right panel). Downregulation of procaspase-3 to 0.1-fold of the baseline value slightly decreased cPARP level at intermediate time points (6 to 12 hours), but did not affect the cPARP level at 24 hours, compared to the baseline model.

**Figure 3.**
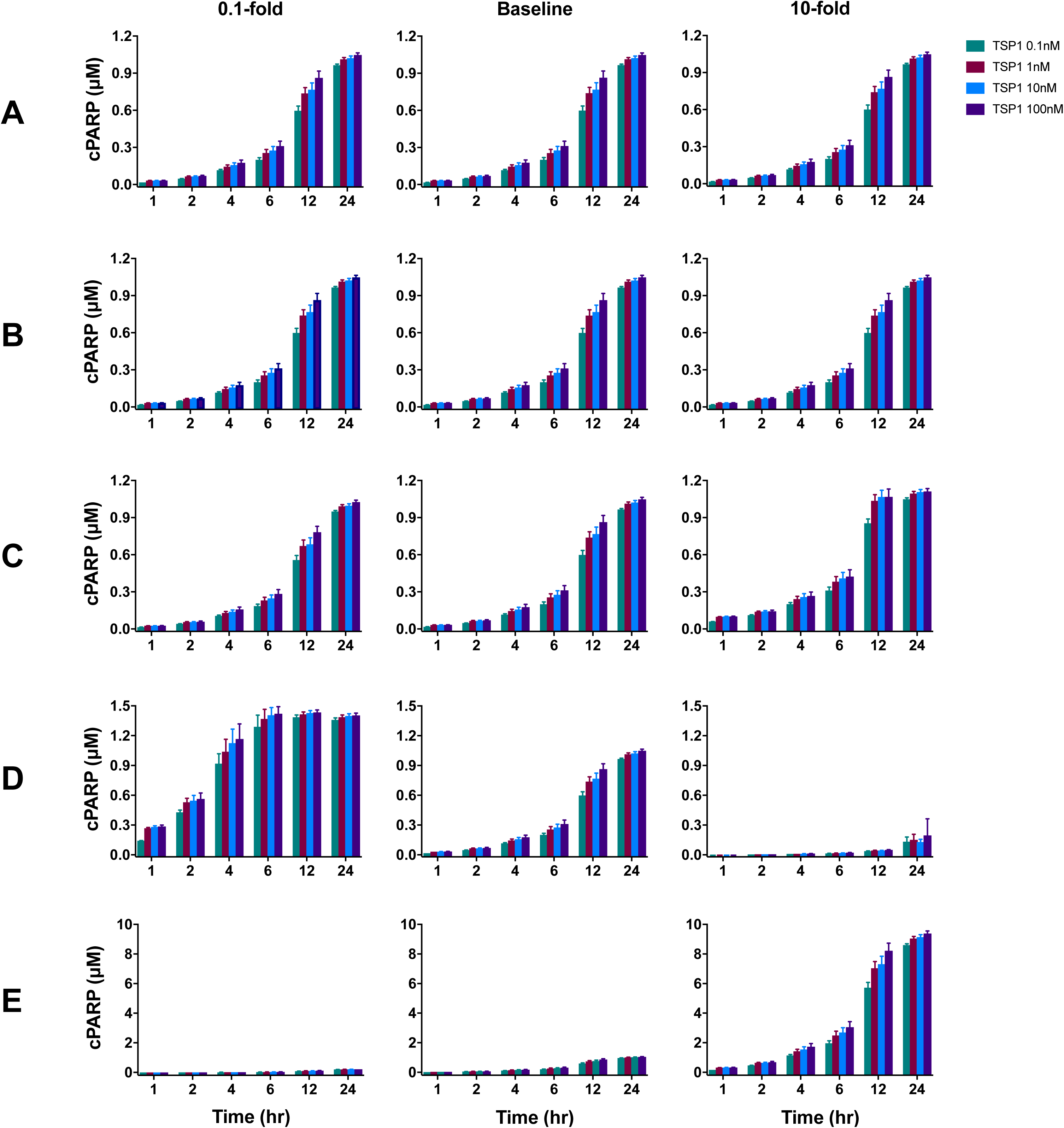
Dose-dependent response of apoptosis signaling with varied initial concentrations. Initial concentrations of A) CD36, B) Fas, C) procaspase-3, D) XIAP, and E) PARP were varied 10-fold above (right column) and below (left column) the baseline values (center column). The model was used to simulate cPARP level in response to four different TSP1 concentrations: 0.1, 1, 10, and 100 nM. The predicted cPARP level at 24 hours was generated using the 12 best sets of parameter values for each condition. The mean cPARP concentration is plotted; error bars show the standard deviation.

Regulation of the caspase-3 inhibitor XIAP reduces apoptosis signaling. That is, increasing the level of XIAP by 10-fold dramatically decreased cPARP concentration to less than 20% of the baseline level, as shown in the right panel of Figure 3D. However, decreasing XIAP by 0.1-fold results in a larger and faster increase in cPARP level compared to the baseline model (Figure 3D, left panel). For example, after 24 hours, the decreased XIAP resulted in 41% and 34% more cPARP than the baseline level, with 0.1 nM and 100 nM TSP1, respectively.

Lastly, the model predicts that increasing PARP levels significantly influences cPARP levels (Figure 3E). When PARP is increased by 10-fold, the cPARP level at all time points is approximately nine times higher than the amount produced in the baseline model. In summary, the apoptotic response stimulated by TSP1 is sensitive to varying the concentrations of certain intracellular species.

### Perturbing the signaling pathway influences the population response to TSP1 stimulation

Next, we implemented perturbations in the model and predicted the response of individual cells in a population. We accounted for heterogeneity in the cell population by introducing randomness in the initial concentrations of protein species. In this population-based model, the initial concentrations of all starting species are drawn from a gamma distribution [41]. Here, the baseline model uses the parameter set that best fit the data (produced the lowest error) out of the 12 parameter sets obtained from model training. We then ran the model 2,000 times, representing 2,000 independent cells, and analyzed the population-level response to TSP1 stimulation. We simulate the response to seven conditions (as described in the Methods) at two TSP1 concentrations (0.1 and 10 nM). Particularly, we investigated whether the predicted results from solving the deterministic model with the fixed initial conditions (Figure 3) hold true when accounting for heterogeneity at the population level.

We characterized the population-level response based on the cells’ cPARP concentration. We extracted predicted intracellular cPARP concentrations for the 2,000 cells at distinct time points up to 24 hours of TSP1 stimulation, and generated histograms. This provides a direct visualization of the distribution of cPARP levels in the cell population. Based on literature data, we defined the threshold of intracellular cPARP required for apoptosis to occur within each cell to be 1.05 μM (see Methods). We used the model to predict the percentage of apoptotic cells, based on the predicted cPARP concentrations. Cells that have high cPARP level (greater than 1.05 μM) are classified as apoptotic, since their cPARP level exceeds the threshold value. Cells whose intracellular cPARP concentration is below the threshold value are classified as non-apoptotic. We also analyzed when cells that have high cPARP level appear at the simulated time points.

In the baseline model, the apoptotic response initiates within six hours after 10 nM TSP1 stimulation (Figure 4A). The size of the cPARP-positive population increases throughout the 24-hour stimulation. By 24 hours, the apoptotic cells make up 41% of the total population (Figure 4E). Below, we compare the population-level response for the baseline model to the response when particular species in the intracellular signaling network are perturbed.

**Figure 4.**
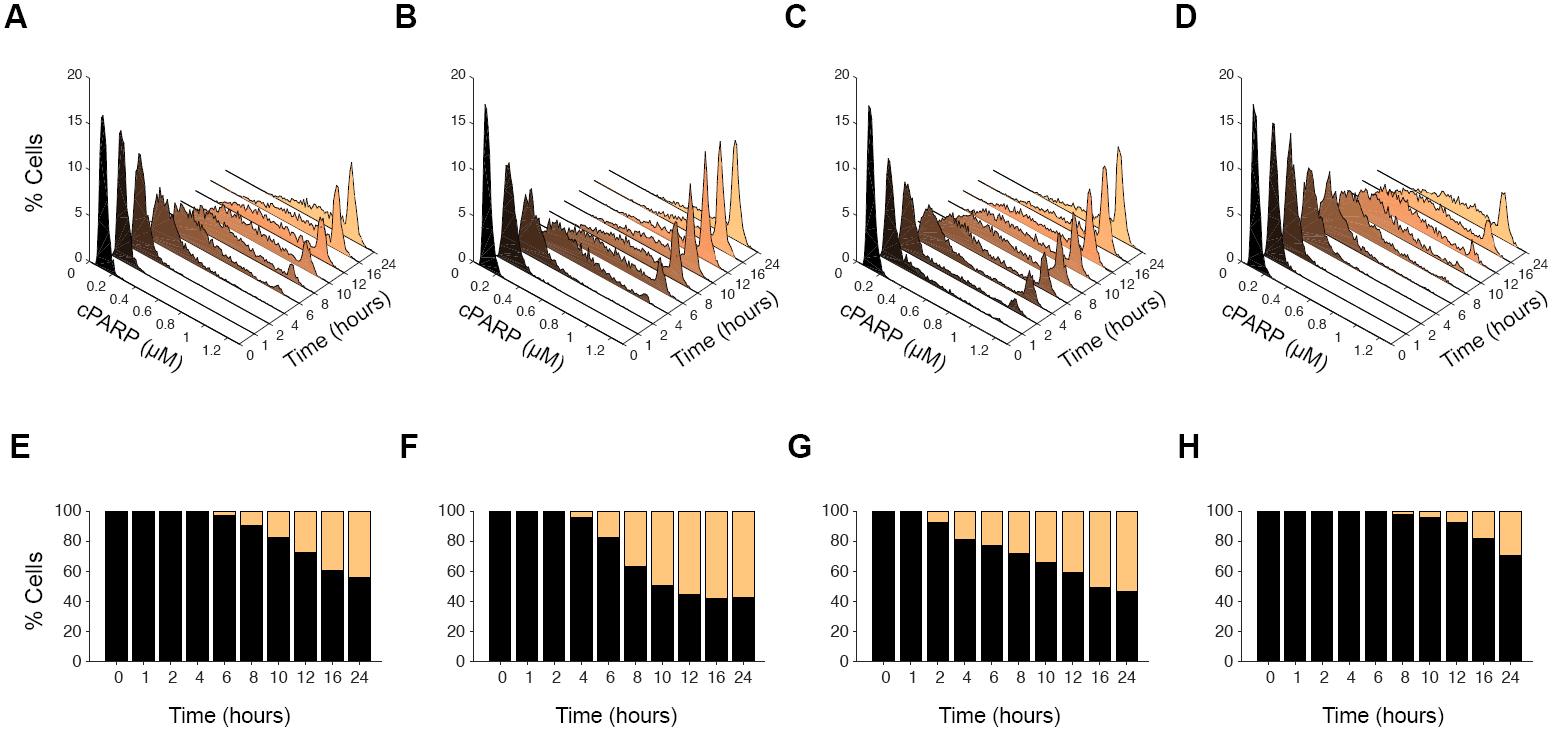
Distribution of cPARP concentration in population-level model. (A)-(D): Histogram showing the percentage of the 2,000 cells with a given cPARP concentration, in response to 10 nM TSP1 stimulation. A) Baseline model; B) XIAP downregulation; C) DXR treatment; and D) Increased nuclear translocation rate. A different color is assigned to each time point. (E)-(H): The predicted percentage of non-apoptotic (black) and apoptotic (orange) cells in response to 10 nM TSP1 stimulation. E) Baseline model; F) XIAP downregulation; G) DXR treatment; and H) Increased nuclear translocation rate.

According to the model with fixed initial conditions, downregulation of XIAP strongly promotes apoptotic signaling (Figure 3D, left panel). To explore whether this conclusion still holds with a heterogeneous cell population, we decreased XIAP expression level by 0.5-fold, a physiologically reasonable change to the protein expression, and simulate the population-based response to 10 nM TSP1 stimulation under this condition. The results show that with XIAP downregulation, cells with high cPARP level appear at four hours (Figure 4B,F). By 24 hours, 57% of the cell population is apoptotic. Additionally, the apoptotic population exceeds the non-apoptotic population by 0.4-fold (Figure 4F).

We also considered the effect of doxorubicin (DXR) on the apoptosis signaling network. A published experimental study suggest that a low dosage of DXR sensitizes cells to pro-apoptotic signaling [32]. Specifically, Quesada and coworkers observed that Fas receptor expression increases approximately 3-fold following DXR treatment. Thus, we simulated the effect of such a DXR treatment by increasing Fas expression by 3-fold. Additionally, we increased the synthesis rates of certain intracellular species, as DXR has been shown to increase protein expression [34]. Here, we increased *K_syn___a_ii* by 10-fold. Under this simulated DXR treatment condition, cells with high cPARP level appear within two hours, a faster apoptotic response than in the baseline model or with XIAP downregulation (Figure 4C). However, the population of cells with high cPARP is 52%, which is not as large as what is predicted with XIAP downregulation (57%). Additionally, with DXR treatment, the progressive increase in the percentage of apoptotic cells through 24 hours is more gradual than with XIAP downregulation (Figure 4F, G). By 24 hours, 52% of the cells are apoptotic, and there are 0.2-fold more apoptotic cells than non-apoptotic cells.

Upon examining structure of the signaling network, we hypothesized that increasing the transiocation rate of phosphoryiated p38MAPK (pp38) and NF-kB into the nucleus can promote apoptotic signaling. Therefore, we simulated the model with K,_rep_ increased by 10-fold. The apoptotic response is slower and smaller in scale than in the baseline model (Figure 4D). Additionally, the positive population does not appear until 10 hours after starting TSP1 stimulation, and by 24 hours, less than 30% of the cells are apoptotic (Figure 4H).

We also simulated the population response with procaspase-3 upregulation, increased Fas expression, phosphatase inhibition, and kinase-activity upregulation. The results are shown in Supplemental Figure S4. These perturbations to the signaling network do not dramatically affect the population response. That is, the speed and magnitude of the response in each case are similar to the baseline model (Figure 4, panels A and E). Apoptotic cells begin to appear by six hours, and after 24 hours of TSP1 stimulation, at least 40% of the cells are apoptotic (Figure S4). Additionally, we predicted the population-based response for the baseline model and the seven network perturbations when the cells are stimulated with 0.1 nM TSP1. These results are shown in Supplemental Figure S5. Next, we present our detailed analysis of the predicted results for the two TSP1 stimulation levels and compare the apoptotic response.

### Strategies to enhance apoptotic response have differential effects on the magnitude and time scale of TSP1-mediated signaling

We applied the model to distinguish the effects of possible strategies to promote TSP1-induced apoptosis under different levels of TSP1 stimulation. Here, we compared three quantities: the percentage of cells that have reached the cPARP threshold by 24 hours, the maximum cPARP level reached in each cell, and the time to reach the threshold cPARP concentration *(T_t_)* within the 24-hour simulation time. Combined with the results shown in Figure 5, these quantities provide insight into TSP1-mediated apoptosis signaling. Below, we compare the results to the baseline case.

**Figure 5.**
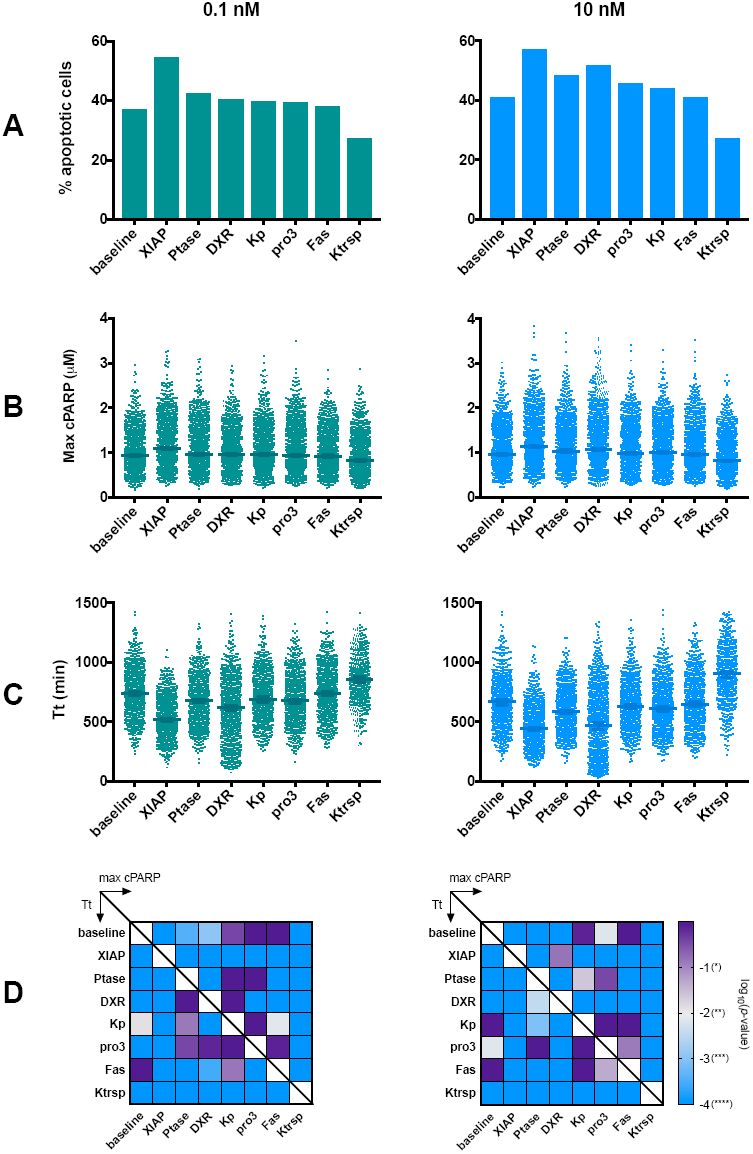
Predicted population-based response to TSP1 stimulation. Comparison of three quantities that characterize the population-level response: A) the percentage of cells that have reached the cPARP threshold by 24 hours, B) the maximum cPARP level reached in each cell, and C) the time to reach threshold *(T_t_).* D) Heat maps representing the statistical analysis comparing the maximum cPARP concentration and *T_t_* in the cell population.

In the baseline model, 0.1 nM TSP1 stimulation leads to 37% of the cells being apoptotic at 24 hours. The model predicts that 49% more cells are apoptotic with XIAP downregulation than the baseline (Figure 5A, left panel). Additionally, phosphatase inhibition and DXR treatment increased the apoptotic population by 14% and 9%, respectively, compared to the baseline model. Upregulation of procaspase-3 or kinase activity both increase the apoptotic population by approximately 7%. Increasing Fas expression did not have an effect on the percentage of apoptotic cells, while increasing the translocation rate decreased the apoptotic cells by 27%.

With 0.1 nM TSP1 stimulation, the median values for the maximum cPARP attained under each simulated condition follow the same order of effectiveness as observed with the percentage of apoptotic cells (Figure 5B, left panel). Additionally, we performed statistical analyses to compare the maximum cPARP between the baseline model and each perturbation (Figure 5D, left panel upper triangle). The maximum cPARP reached is highly significantly different from the baseline model when XIAP is downregulated (the maximum cPARP is higher; p<0.0001) or when the translocation rate is increased (cPARP decreases; p<0.0001). There are also significant differences in the predicted maximum cPARP when comparing the perturbations (Figure 5D, left panel, upper triangle). Notably, the effects of altering XIAP or the nuclear translocation rate are significantly different from all of the other strategies.

Interestingly, the effectiveness of the approaches on time to reach threshold does not follow the same order as that of the percentage of apoptotic cells or of maximum cPARP (Figure 5C, left panel). Based on our statistical analysis, compared to the baseline level, the time to reach threshold is significantly shorter (with high significance level) with XIAP downregulation, phosphatase inhibition, DXR treatment or procaspase-3 over-expression (Figure 5D, left panel, lower triangle). In contrast, the time to reach threshold is significantly longer when the translocation rate is increased by 10-fold.

With 10 nM TSP1 stimulation, XIAP downregulation leads to 40% more apoptotic cells than the baseline model (Figure 5A, right panel). Phosphatase inhibition and DXR treatment also lead to a strong response, with 17% and 27% more apoptotic cells, respectively. Increasing procaspase-3 or kinase activity both increased the relative size of the apoptotic population by approximately 10%, compared to the baseline.

XIAP downregulation, phosphatase inhibition, DXR treatment, and increase of procaspase-3 expression significantly increased the maximum cPARP level and shortened the time to reach threshold (Figure 5B-D, right panels). Increasing the translocation rate significantly decreased the maximum cPARP reached, and prolonged the time to reach threshold.

In summary, these results show that XIAP downregulation is more effective than the other approaches in both increasing the maximum cPARP level attained in the cell population and shortening the time to reach the cPARP threshold. This holds true when cells are stimulated with either 0.1 nM or 10 nM TSP1. Increasing the level of TSP1 stimulation to 10 nM makes all of the pro-apoptotic strategies more effective. However, it is interesting to note that the ordering of the strategies from most effective to least effective changes with the level of TSP1 stimulation.

### Initial protein expression levels influence apoptosis response

In order to explore the cause for the different responses to the apoptosis signaling within the population of 2,000 cells, we compared the initial conditions of the apoptotic cells and non-apoptotic cells at 24 hours. Statistical analysis of the distributions of the initial protein concentrations (normalized to their baseline values) indicate that XIAP concentration in the apoptotic population is significantly lower than in the non-apoptotic population (p<0.0001), while PARP concentration is significantly higher in the apoptotic population (p<0.0001). In fact, the relationship between the apoptotic response and the XIAP and PARP initial conditions is easily visualized (Figure 6A). This illustrates that apoptotic cells (Figure 6A, orange markers) have higher PARP and lower XIAP than non-apoptotic cells (Figure 6A, black markers). The NF-kB concentration in the cytosolic compartment is also significantly lower in the apoptotic population compared to apoptotic cells (p=0.04). Thus, statistical analysis shows that the initial concentrations of certain species distinguish apoptotic from non-apoptotic cells.

**Figure 6.**
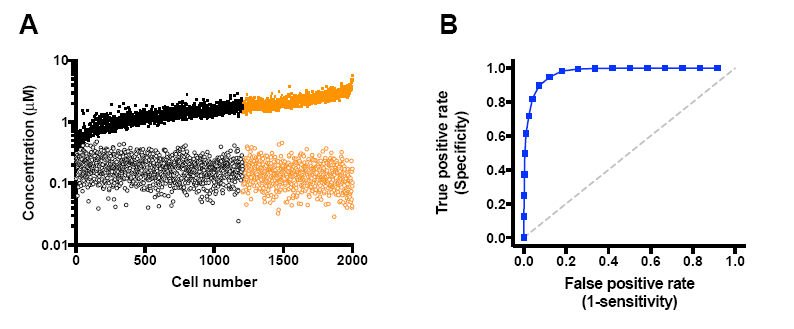
Figure 6. Relationship between initial conditions and predicted apoptotic response. A) Initial concentrations of PARP (squares) and XIAP (circles) for apoptotic cells (orange) and non-apoptotic cells (black). The cells are ordered according to the maximum cPARP attained within the 24-hour simulated TSP1 stimulation. The difference between the initial amounts of PARP and XIAP is higher for apoptotic cells, which reach higher cPARP levels. B) ROC curve for the difference between the initial concentration of PARP and XIAP input. The AUC is 0.97 and is significantly difference than 0.5 (given by the gray dashed line), indicating that this input is a reliable predictor of the cells’ response to TSP1 stimulation.

Building on the results from the statistical analysis of the initial concentrations, we investigated whether the initial concentrations can be used to classify cells as apoptotic or non-apoptotic upon 24 hours of TSP1 stimulation. Here, we generated a receiver operator characteristic (ROC) curve. The ROC curve illustrates the ability of an input descriptor to classify a response with high sensitivity and specificity (see Methods). For inputs, we used the initial concentration of each species that does not start at zero (16 species), the ratios of XIAP and PARP concentrations ([XIAP]/[PARP] and [PARP]/[XIAP]), and the absolute value of the difference between the concentrations of PARP and XIAP] (i.e., |[PARP]-[XIAP]|). Thus, we considered 19 total inputs that may possibly predict the response to TSP1-mediated apoptosis. We use the area under the ROC curve (AUC) to compare the ability of the inputs to predict the response to TSP1.

Constructing the ROC curve for the 19 inputs shows that the absolute difference between the initial concentrations of XIAP and PARP predicts which cells will be apoptotic. Specifically, the difference between XIAP and PARP can classify the cells with high sensitivity and specificity (Figure 6B). In this case, the AUC is 0.97, with a 95% confidence interval of [0.96 – 0.98], providing a quantitative measure of the predictive ability of the difference between the XIAP and PARP concentrations. This AUC is significantly different than 0.5 (p<0.0001), which indicates that using the difference between the concentrations of XIAP and PARP is more predictive than classifying cells as apoptotic or not based purely on chance. The AUC values for classifying the apoptotic response using the initial concentration of XIAP or PARP alone are 0.65 and 0.95, respectively, and in both cases, the AUC values are significantly different from 0.5 (p<0.0001). Thus, although using the concentration of XIAP or PARP alone is better than randomly guessing which cells will respond, these concentrations are less reliable predictors when considered individually. The AUC values for the remaining 16 inputs range from 0.48 to 0.54, and are not significantly different than randomly selecting which cells will be apoptotic. Quantitative results from constructing the ROC curve for all of the inputs are provided in Supplementary Table S5. Overall, the results of this analysis demonstrate that the initial concentrations of PARP and XIAP, and especially the difference between their concentrations, can be used to predict which cells will respond to TSP1 signaling.

## Discussion

We have developed a molecular-detailed model of the TSP1-induced apoptotic signaling. The model represents the reaction network of interactions involved in the intracellular signaling pathway and includes multiple feedback loops. Our model captures the feedback from transcriptional regulation by NF-kB onto Fas signaling, which allows us to expand the dynamics of the network to a longer time scale. Predictions from the trained model match experimental data. We further validated the model using a separate set of experimental measurements. We implement the model using ODEs, which can be solved once to simulate the average response of a population of cells. We also account for randomness in the protein concentrations by solving the ODEs 2,000 times with varied initial conditions to predict the individual responses of 2,000 cells. In this case, the cells each have different initial concentrations of the molecular species, representing heterogeneity of the cell population. This heterogeneity influences the responses to TSP1 treatment and the effectiveness of strategies aiming to increase apoptosis signaling.

The model provides mechanistic and quantitative explanations of the effects of several approaches to promote TSP1-mediated apoptosis. Using the model, we proposed and simulated the effects of perturbing the signaling network, including altering receptor and protein levels, rates of protein synthesis and transport, the activity of specific phosphatases, and the overall kinase activity within the cells. These simulations exploit the power of mathematical modeling to generate quantitative predictions that would otherwise be time‐ and cost-consuming to explore experimentally. Overall, our model provides quantitative insight into the effects of targeting particular aspects of the TSP1-mediated apoptosis signaling pathways.

The model predicts that downregulation of XIAP is a promising strategy to enhance TSP1’s apoptotic signaling effect, sensitizing cells to low-dosage TSP1 treatment. XIAP is a potent inhibitor of caspase activity [33]. Experimental studies have shown that specific over-expression of XIAP can rescue cells from apoptosis, and antisense downregulation of XIAP led to a dramatic decrease in resistance to radiation-induced cell death [42]. Our model simulation shows that in TSP1-mediated apoptotic signaling, modulating XIAP level also has a similar effect. Importantly, the model provides quantitative and mechanistic insight into the effects of targeting XIAP. Downregulation of XIAP directly leads to an increased level of active caspase-3, a crucial mediator in the apoptosis signaling network. Our simulation results demonstrated that this approach is the most effective in promoting endothelial cell apoptosis. The further analysis based on the ROC curve supports the model simulations showing the importance of XIAP in influencing the dynamics of TSP1-mediated apoptosis. Interestingly, the ROC curve reveals a relationship between XIAP and PARP that we had not identified using model simulations alone. The difference between the initial concentrations of those proteins can accurately predict which cells will undergo apoptosis in response to TSP1 stimulation, even before the cells are exposed to TSP1.

In another example, a strategy to increase apoptosis signaling is supported by experimental studies. We used the model to quantify the effect of inhibiting MAPK phosphatase (MKP) activity. We simulated the effect of this approach by decreasing the binding affinity between the phosphatase and its substrate, pp38, and the dephosphorylation rate. Therefore, the active pp38 level remains high as the phosphatase activity is inhibited, enabling downstream signaling. The results indicate that this approach can effectively promote the apoptotic response. Our predictions agree with experimental results that show that modulating MKPs is a viable option to promote apoptosis mediated by p38MAPK [28,29].

The model also generates non-intuitive results. The simulations show that increasing the rate of translocation from the cytosol to the nucleus delayed and attenuated the apoptotic response. This observation is counterintuitive, as faster translocation of species immediately upstream of FasL production is expected to accelerate signal transduction. However, the model simulation suggests that with faster translocation, the pool of caspase-3 and p38MAPK is depleted in upstream signal transduction, before DISC formation occurs (data not shown). The effect of a pan-kinase activity promoter is another example of non-intuitive predictions. We simulated the kinase promoter by increasing the phosphorylation rates of p59fyn, p38MAPK, and IkB by 10-fold. Intriguingly, this approach did not affect either the response time (time to reach apoptosis threshold) or the percentage of apoptotic cells. The explanation is that increased kinase activity depleted certain species before their accumulation is achieved, in a similar manner to the faster cytosol-to-nucleus translocation case.

In addition to proposing strategies to increase apoptosis signaling, the model provides mechanistic insight into experimental observations. At low dosages, doxorubicin treatment has been shown to promote TSP1-mediated apoptosis. Using our model, we propose the potential mechanism of action of this effect. A study by Quesada *et al.* showed that endothelial cell apoptosis was due to a synergistic effect of the upregulation of FasL induced by TSP1 and upregulation of Fas by doxorubicin [32]. Our model simulations show that altering Fas receptor expression level alone does not affect the apoptotic response; rather, the availability of intracellular signaling species significantly influences apoptosis signaling as well. Specifically, the model simulations show that increasing the synthesis of particular molecular species is required to qualitatively match the effects of DXR observed experimentally. Based on these modeling results, we hypothesize that the low dosage doxorubicin treatment not only regulates the protein expression of Fas, but also other species. Our predictions agree with other experimental studies that show translation of multiple proteins increases when the cells are exposed to stress [25,30]. Our model simulation demonstrates the effect of a low dosage of doxorubicin through the proposed mechanism of action, which can be further tested with experiments.

This work provides a biophysically realistic network that generates reliable predictions of the population-level responses. We incorporated stochasticity into the ODE model to represent a heterogeneous population of cells. This implementation provides a framework to test how molecular-targeted strategies influence individual cells in a population. Solving the ODE model only once for a single set of initial conditions indicates that upregulation of procaspase-3 greatly increases the magnitude of the apoptotic response, while procaspase-3 downregulation does not affect cPARP level. This implies that a threshold value of procaspase-3 expression may be needed in order to enhance apoptosis signaling. On the other hand, the population-based model simulations showed that overexpressing procaspase-3 by 3-fold has only a mild effect of increasing the apoptotic response. These conflicting results demonstrate the importance of accounting for heterogeneity within a cell population. Therefore, we emphasize the utility of the population-based model to make predictions, as the deterministic model that represents the dynamics of an average single cell can be misleading in certain cases.

To our knowledge, this is the first mechanistic model to investigate TSP1-mediated apoptosis. The apoptotic signaling in this study is essential to the inhibitory effect of TSP1 [15]. Notably, TSP1 not only induces apoptosis through the CD36 receptor, but also has many other anti-angiogenic functions, possibly making it more potent than a single apoptosis inducer for regulation of angiogenesis. The framework established in our study can be readily adapted and combined with models describing other interactions of TSP1 [44], or pro-angiogenic signaling networks [22,23] for future modeling studies. The model can also be expanded to include cell-cell interactions with both exogenous and endogenous FasL signaling. Future work can improve the model framework in various ways, for example, by adding the downstream function of PARP to address the balance between survival and apoptotic effects as PARP loses its repair function in its cleavage; specifying initial concentrations from different cell types; including more detailed reaction mechanisms (such as NF-kB activation by p38MAPK); or accounting for the mitochondrial feed-forward loop, which provides the link to the intrinsic apoptotic pathway.

Our model complements other studies that focus on apoptosis signaling promoted by death ligands and their receptors [32–34]. We adapted the model for Fas-mediated apoptotic and NF-kB signaling established by Neumann *et al.,* which served as a foundation for the FasL signaling cascade in our work. In other work, Albeck *et al.* established a model to describe TNF-related apoptosis-inducing ligand (TRAIL) induced apoptosis in HeLa cells (an ovarian cancer cell line), with a focus on the “variable-delay, snap-action” switching mechanism of extrinsic apoptosis. Both the Albeck model and our model exhibit the cells’ response of switching to a high apoptotic response. However, one key difference between our model and theirs is how the switching arises. In the Albeck model, all-or-none switching of the cell death response is achieved by a network that does not include feedback. They concluded that this snap-action arises from interplay between the biochemistry of protein-protein interaction and translocation between the cytosol and mitochondria. We have simplified this extrinsic apoptosis pathway in our model; however, the snap-action behavior of apoptosis is still evident, shown by the fast accumulation of cPARP. This switching is a direct result of the reaction between caspase-3 and PARP, amplified by the feedback loops. Another difference between the two models is how stochasticity is implemented. The time to reach threshold *(T_t_)* in our model is analogous to the delay time *(T_d_)* in Albeck model. Albeck and co-workers impose a distribution in the delay time by randomly selecting *T_d_* from a defined range. In contrast, in our model, cell-to-cell differences in the switching delay emerge solely based on the variation in intracellular protein concentrations. This is a more realistic framework that represents an actual population of cells. Thus, our model is a tool to analyze the population-level responses to TSP1 stimulation and perturbations to the signaling network.

## Conclusions

In summary, our model quantitatively describes the TSP1-mediated intracellular signaling via the CD36 receptor, which leads to endothelial cell apoptosis. This model predicts that downregulation of XIAP is the most promising way to effectively promote TSP1-mediated apoptosis. In addition, we propose an alternative mechanism of action for the effect of low dosage doxorubicin treatment in sensitizing cells to TSP1 stimulation. This model framework can be ultimately used to generate and optimize TSP1-based therapeutic strategies for promoting apoptosis.

## Acknowledgements

The authors thank members of the Finley research group for critical comments and suggestions and Dr. James Finley for insight regarding the ROC analysis.

## Funding

The authors acknowledge the support of the US National Science Foundation (CAREER Award 1552065). The funders had no role in study design, data collection and analysis, decision to publish, or preparation of the manuscript.

## Supplemental Materials

**List of Supplemental Tables (see File S1.xlsx)**

Table S1. Biochemical reactions and parameter values

Table S2. Initial protein concentrations

Table S3. Estimated parameter values

Table S4. Results from ANOVA analysis for *T*_*t*_ and max cPARP

Table S5. Results from ROC analysis

**Model Equation (see File S2.pdf)**

Ordinary differential equations and initial conditions

**List of Supplementary Figures (see File S3.pdf)**

Figure S1. Comparison of minimal model of FasL signaling to experimental data

Figure S2. Global sensitivity analysis

Figure S3. Distribution of estimated parameter values

Figure S4. Population-level response to TSP1 stimulation

Figure S5. Population-level response to 0.1 nM TSP1 stimulation

